# Geographical variability of bacterial communities of cryoconite holes of Andean glaciers

**DOI:** 10.1101/2021.01.14.426633

**Authors:** Francesca Pittino, Michael Seeger, Roberto Azzoni, Roberto Ambrosini, Andrea Franzetti

**Affiliations:** Department of Earth and Environmental Science, University of Milano-Bicocca, Milano, Italy; Departamento de Química & Centro de Biotecnología (CBDAL), Universidad Técnica Federico Santa María, Valparaíso, Chile; Department of Environmental Science and Policy, University of Milan, Milano, Italy; Department of Earth Sciences “Ardito Desio”, Milano, Italy

## Abstract

Cryoconite holes, ponds full of melting water with a sediment on the bottom, are hotspot of biodiversity of glacier surface. They host a metabolically active bacterial community that is involved in different dynamics concerning glacier ecosystems. Indeed, they are responsible of organic matter production and with other microorganisms establish a real microecosystem. Cryoconite holes have been described in different areas of the world (e.g., Arctic, Antarctic, Alps, Himalaya), and with this study we will provide the first description of bacterial communities of cryoconite holes of the Andes in South America. We collected samples on three high elevation glaciers of the Andes (Iver, Iver East and Morado glaciers) and two Patagonian glaciers located at sea level (Exploradores glacier and Perito Moreno). Results show that the most abundant orders are Burkholderiales, Cytophagales, Sphingobacteriales, Actinomycetales, Pseudomonadales, Rhodospiarillales, Rhizobiales, Sphingomonadales and Bacteroidales, which have been reported on glaciers of other areas of the world, Bacterial communities change from one glacier to another and both water pH and O2 concentration affect bacterial communities composition.

## Introduction

Glaciers and ice sheets have been recognized as a biome in their own right as they host viable populations of organisms (Anesio and Laybourn-Parry 2012). Glacier ecosystems are mostly dominated by microorganisms that have been found in all parts of the glacier environment, including the englacial and the subglacial zones (Anesio and Laybourn-Parry 2012). The supraglacial zone is the most biodiverse among glacier ecosystems (Hodson et al. 2008) and, for practical reasons, is also the most studied. Here, small depressions of the glacier surface filled by meltwater, the cryoconite holes, form as a consequence of the atmospheric deposition of a fine-grained sediment, cryoconite, that locally decreases the albedo and melts the underlying ice. The main source of this sediment is the surrounding environment, but a small part of it is deposited after long-range atmospheric transport (Franzetti et al. 2017a). In the extreme glacier environment, cryoconite holes are protected habitats where general conditions are more favourable for life than in the other parts of a glacier, as they provide liquid water and act as a protective barrier against the intense UV radiation characteristic of the glacier environment (McIntyre 1984). At low temperature, an ice lid can form on the water surface in a cryoconite hole, which further protects this microhabitat. In fact, underneath this cover, water remains liquid because solar radiation can penetrate the ice lid and be absorbed by the dark cryoconite; only in case of extremely cold temperatures the water inside the hole completely freezes (Fountain et al. 2004a).

Cryoconite holes are present on glaciers in almost all areas of the world; so far they have been described in the Arctic, Antarctica, on the Alps, in Tien Shan and in Himalaya (Takeuchi, Nishiyama, and Li 2001;Mueller and Pollard 2004; Telling et al. 2010; Uetake et al. 2016; Franzetti et al. 2017a). However, they do not show the same features in all the different geographic areas where they occur. Polar cryoconite holes, in particular, are more stable than those that form on temperate mountain glaciers. On Himalayan glaciers they can sometimes survive more than one year, but this does not always occur (Takeuchi et al. 2000; Takeuchi, Nishiyama, and Li 2010). Indeed, in most temperate mountain glaciers they are mostly ephemeral structures that can be destroyed and form again because of the strong ablation that can quickly dismantle them and wash away the sediment and the intense solar radiation that can form new holes in a few days (Fountain et al. 2004a; Cook, Edwards, Takeuchi, et al. 2016; Pittino et al. 2018).

Communities inhabiting these microhabitats also differ between polar and temperate mountain cryoconite holes. In polar environments bacterial communities in cryoconite holes are quite stable (Musilova et al. 2015), while on the Alps, the only temperate mountain range where ecological successions studies have been conducted on cryoconite holes, they change along the ablation season (Franzetti et al. 2017b; Pittino et al. 2018). In particular, filamentous Cyanobacteria and other phototrophs are the first colonizers of cryoconite holes after their formation at the beginning of the ablation season, while at its end bacterial communities are mostly heterotrophic (Franzetti et al. 2017b). Such a difference between stable polar and seasonally varying temperate cryoconite hole communities may be due to the more stable temperatures along the ablation season in the polar regions. Indeed Pittino et al. (2018) reported an effect of temperature variation along the ablation season on bacterial communities composition on an Alpine glacier.

However, investigations of the temporal variability of cryoconite hole bacterial communities of the same glacier along one or even more ablation seasons were seldom conducted, mostly because of the difficulties and the costs of visiting repeatedly glaciers, which usually occur in remote areas. Most often, the description of the bacterial communities of cryoconite holes is conducted with snapshot studies (Edwards et al. 2013; Stibal, Šabacká, and Kaštovská 2006; Hodson et al. 2010), which allowed a general description of their biotic communities, even if they cannot include the whole biodiversity of these environments (Franzetti et al. 2017b; Pittino et al. 2018). These studies showed that filamentous Cyanobacteria are the main responsible of cryoconite grain formation (Takeuchi, Nishiyama, and Li 2001; Stauch-White et al. 2017; Uetake et al. 2016) and the other most abundant bacterial phyla found in these microhabitats are Proteobacteria (Alfa and Beta), Actinobacteria, Chloroflexi, Acidobacteria and Bacteroidetes (Boetius et al. 2015; Liu et al. 2017; Christner, Kvitko, and Reeve 2003; Franzetti et al. 2016). At order level instead we can mention: Sphingobacteriales, Pseudomonadales, Rhodospirillales, Burkholderiales and Clostridiales (Andrea Franzetti et al. 2017b; Łukasz Kaczmarek et al. 2015). So far, snapshot studies have been used to describe bacterial communities inhabiting cryoconite holes on glaciers in the Alps (Edwards et al. 2013; Franzetti et al. 2017a), Greenland (Musilova et al. 2015; Uetake et al. 2016), the Arctic (Singh, Singh, and Dhakephalkar 2014; Stibal, Šabacká, and Kaštovská 2006), Antarctica (Sommers et al. 2018; Christner, Kvitko, and Reeve 2003), Himalaya (Sanyal et al. 2018), Karakoram (Ambrosini et al. 2017) and China (Takeuchi, Nishiyama, and Li 2001).

To the best of our knowledge, no published information is currently available about cryoconite bacterial communities from South American glaciers. Only one study by Tacheuki et al. (2001) described cryoconite characteristics of the Tyndall Glacier, but from a physico-chemical point of view, not reporting bacterial communities composition. A study reported some data about bacterial communities of glaciers surface describing the gut microbiome of the glacier stonefly (*Andiperla willinki*) that is likely to feed on supraglacial bacteria (Murakami et al. 2018), but no more studies report bacteria inhabiting this environment in South America.

The aim of this work is to provide a first description and an evaluation of the geographical variability of bacterial communities of cryoconite holes of South America, based on samples collected on four Chilean and one Argentinian glaciers. Cryoconite samples were collected from five different glacier of the South American Continent (Fig.1). Three glaciers are located on the Chilean Andes close to Santiago de Chile (Morado, Iver and East Iver glaciers). Two other glaciers (Exploradores Glacier in Chilean Patagonia and Perito Moreno in Argentinian Patagonia) are extremely different compared to the first three mainly due to the different climate setting of the area where are located (Pfeffer et al. 2014).

**Figure 1.**
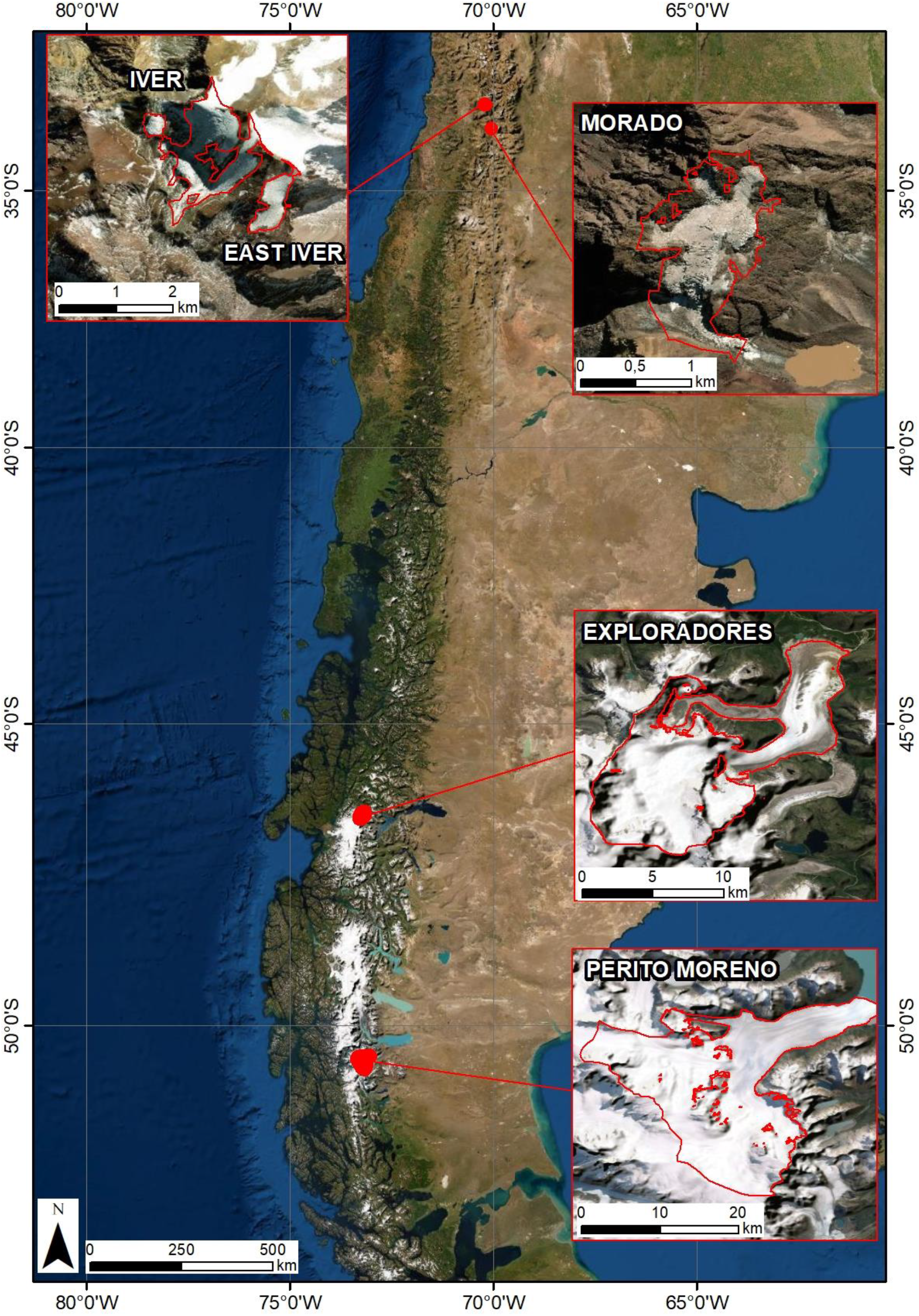
Location Map of glaciers considered in this study. Glacier limit and morphological parameters are obtained from Randolph Glacier Inventory 6.0 (RGI Consortium 2017).

Despite this study was based on snapshot sampling of five glaciers, we are confident that our results can help filling this gap of knowledge on the glacier biodiversity of South America.

## Materials and Methods

### Study areas

Morado Glacier is a valley glacier located about 60 kilometres South East from Santiago, Chile. It covers 1.1 km^2^ and its altitudinal range is between 4604 m a.s.l. and 3535 m a.s.l. It is a small glacier, highly crevassed and partly debris-covered in the ablation area, terminating in a small lake.

Iver Glacier can be classified as a mountain ice body and is located about 30 kilometres North West from Santiago, Chile. It has a surface area similar to Morado, covering 1.64 km^2^ and its altitudinal range is between 5358 m a.s.l. and 4296 m a.s.l. It is a steep glacier, partly covered in the ablation area. During the Little Ice Age, this glacier was probably connected to East Iver Glacier that is a small glacier covering 0.3 km^2^ located few hundred meters from the Iver Glacier.

Exploradores Glacier is a valley glacier located in the Chilean Patagonia. It covers 85 km^2^, it is 19 km long and its altitudinal range is between 3735 m a.s.l. and 158 m a.s.l. The frontal area of the ablation zone is debris-covered.

Perito Moreno Glacier is a valley glacier located in the Argentinian Patagonia. It covers 263 km^2^, it is more than 32 km long and its altitudinal range is between 2800 m a.s.l. and 190 m a.s.l. where the glacier snout ends in a lake, losing most of its mass through calving processes.

### Sample collection

Cryoconite samples were aseptically collected in falcon tubes on five glaciers in 2017 and 2018 (Table 1). We collected samples from 15 cryoconite holes per glacier and, at each hole, we also recorded dissolved O_2_ concentration and pH with a portable oximeter/pH meter (HACH LANGE HQ40D, Loveland, CO, USA).

### DNA extraction and sequencing

DNA was extracted from 0.7 g of cryoconite with the FastDNA® Spin for Soil kit (MP Biomedicals, Solon, OH, USA) according to the manufacturer’s instructions. DNA sequencing was performed on the V5-V6 hypervariable region of the 16S rRNA gene as previously described in Pittino et al. (2018). Sequences were then demultiplexed according to the indexes and clustered in Amplicon Sequences Variants (ASVs) with DADA2 (Callahan et al. 2016). ASVs were then taxonomically classified using rdp classifier (Q. Wang et al. 2007) keeping the full classification only of the taxa attributed with a confidence of 0.8 or higher. Cyanobacteria were classified at phylum level only as rdp classifier does not provide the full classification for them (Garrity et al. 2007; Wilmotte and Herdman 2015).

### Statistical analyses

Analyses were performed with R 3.5.1 (R Core Team, 2014) with the VEGAN, BIODIVERSITYR, MULTTEST, and MULTCOMP packages. Singletons (ASVs present once in one sample only) were removed because they can inflate the variance explained by multivariate tests (Borcard, Gillet, and Legendre 2011) Alpha-diversity was measured using the Shannon diversity index, which accounts for both the richness and the evenness of the species (Shannon 1948), and the Gini inequality index, which is an index of inhomogeneity largely used in economics (Gini 1912). Gini index ranges from 0 to 1 and low values indicate an homogeneous distribution of the species in the community, while high values indicate an heterogeneous distribution (Gini 1912).

Beta diversity analyses were based on the Hellinger distance, which depends on the differences in the ASV proportion among samples, decreases the importance of ASV abundance over occurrence and avoids the double-zero problem when comparing ASV composition among samples (De Cáceres, Legendre, and Moretti 2010; Pierre Legendre and Gallagher 2001). We performed a redundancy analysis (RDA) and a variation partitioning (VP) to quantify the variation of community structures among glaciers (five-level factor), and according to dissolved oxygen concentration and pH of the water above the sediment. The last two variables were mean-centred within glacier before the analyses (i.e. we subtracted from the values recorded at each glacier their mean value). Hereafter, centred variables will be called ΔpH and Δ[O_2_], respectively. Post-hoc tests were also performed to assess pairwise differences between glaciers while correcting P-values for multiple testing according to the false discovery rate (FDR) procedure (Yoav Benjamini and Yekutieli 2001). Variation in the abundance of the 9 most abundant orders according to the variables that significantly affected the structure of bacterial communities identified by the RDAs was investigated by generalized linear models (GLMs) assuming a Poisson distribution and correcting for overdispersion. Also in these cases, P-values were corrected using the same FDR procedure as above. Generalized least-square (GLS) models accounting for heterogeneity of variance among glaciers were also used to investigate changes in oxygen concentration, pH and alpha diversity indices according to the same predictors. No deviation from normality assumption was detected during routine model checks (details not shown).

## Results

Both pH (F_4,70_ = 11.87, P = < 0.001) and oxygen concentration (F_4,70_ = 281.9, P < 0.01) in cryoconite holes differed significantly among glaciers. Post-hoc tests highlighted that glaciers could be divided into two groups with respect of pH values, with Exploradores and Morado showing higher values than East Iver and Perito Moreno (|t_70_| ≥ 0.147, P ≤ 0.035; Fig .2a), while all glaciers differed to one another in their oxygen concentration (|t_70_| ≥ 3.480, P ≤ 0.011; Fig. 2b).

**Figure 2.**
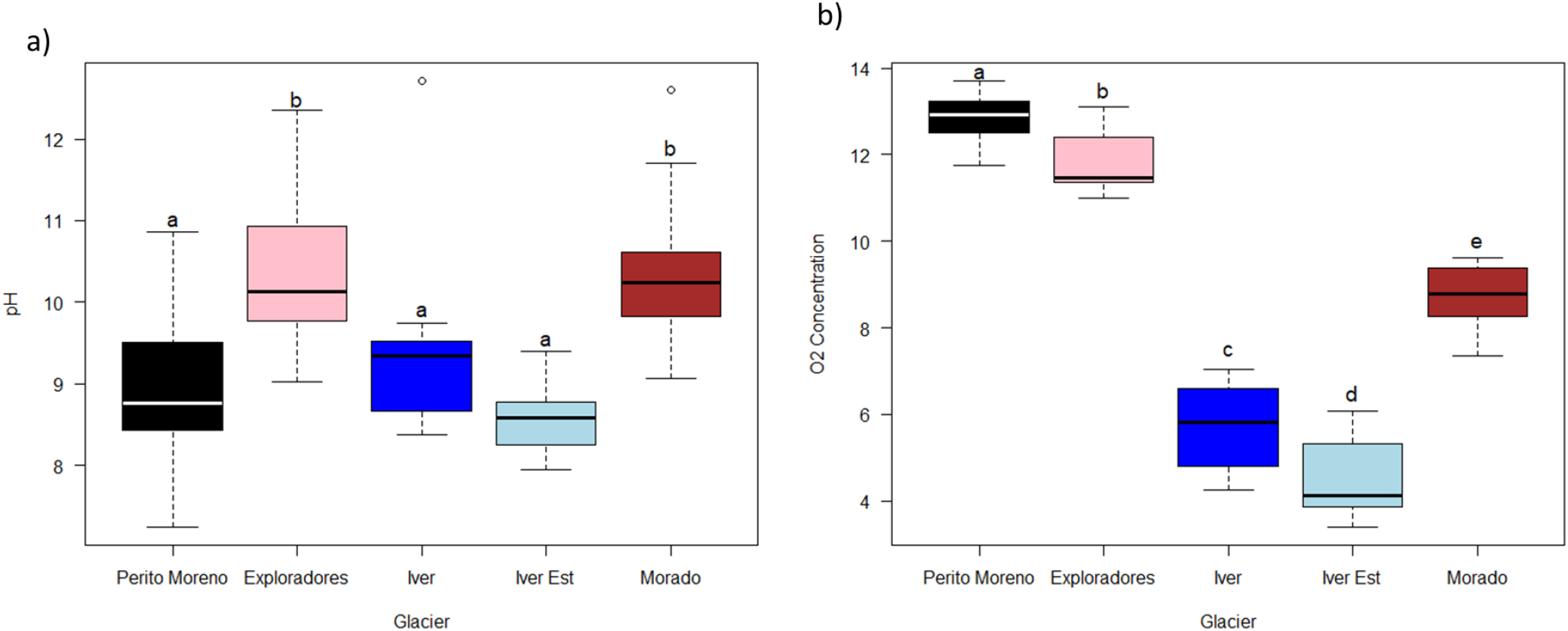
Boxplots of pH (a) and oxygen concentration (b) of cryoconite holes in the investigated South American glaciers. The thick lines represent the median, boxes upper and lower limits the 25^th^ and the 75^th^ percentiles respectively, whiskers the data that go beyond the 5^th^ and the 75^th^ percentile, dots represent the outliers and different letters indicate significant differences between glaciers at post-hoc tests.

We obtained 7,600 − 93,425 sequences per sample. Orders with more than 24,000 sequences were considered as the most abundant. They were: *Burkholderiales, Cytophagales, Sphingobacteriales, Actinomycetales, Pseudomonadales, Rhodospiarillales, Rhizobiales, Sphingomonadales* and *Bacteroidales*.

The RDA performed on all the samples, showed that bacterial communities differed significantly among glaciers and varied according to ΔpH and Δ[O_2_] (Table 2; Fig. 4). The biplot also showed that cryoconite hole bacterial communities of the three small glaciers in central Chilean Andes (Iver, East Iver and Morado) seem to vary mostly according to oxygen concentration, while those of the Exploradores glacier seem to be affected by pH.

**Table 2.** Date of sampling of each glacier.

| Glacier | Date |
| --- | --- |
| Morado | 18/02/2018 |
| Iver | 21/02/2018 |
| East Iver | 22/02/2018 |
| Exploradores | 01/03/2018 |
| Perito Moreno | 29-30/03/2017 |

**Table 3.**
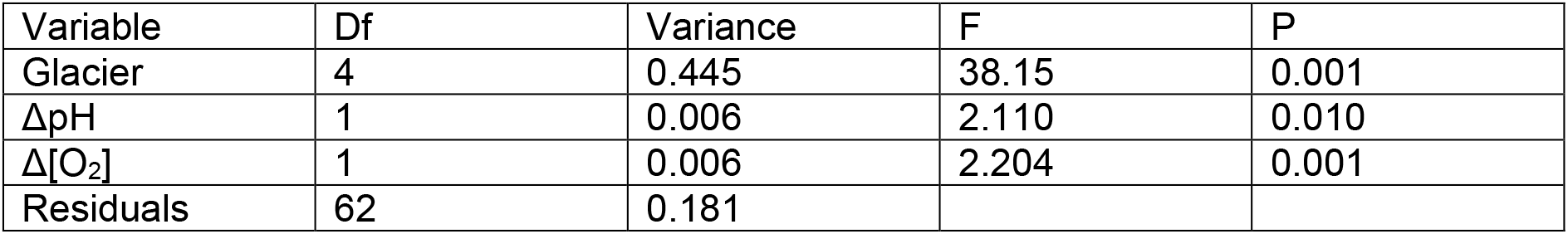
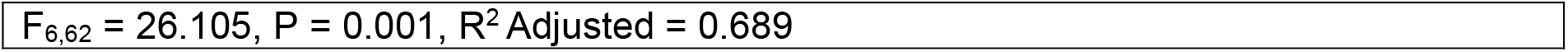
RDA of Hellinger-transformed bacterial ASVs abundances of all the samples according to the glacier, pH and oxygen concentration centred according to their mean value per glacier.

**Figure 3.**
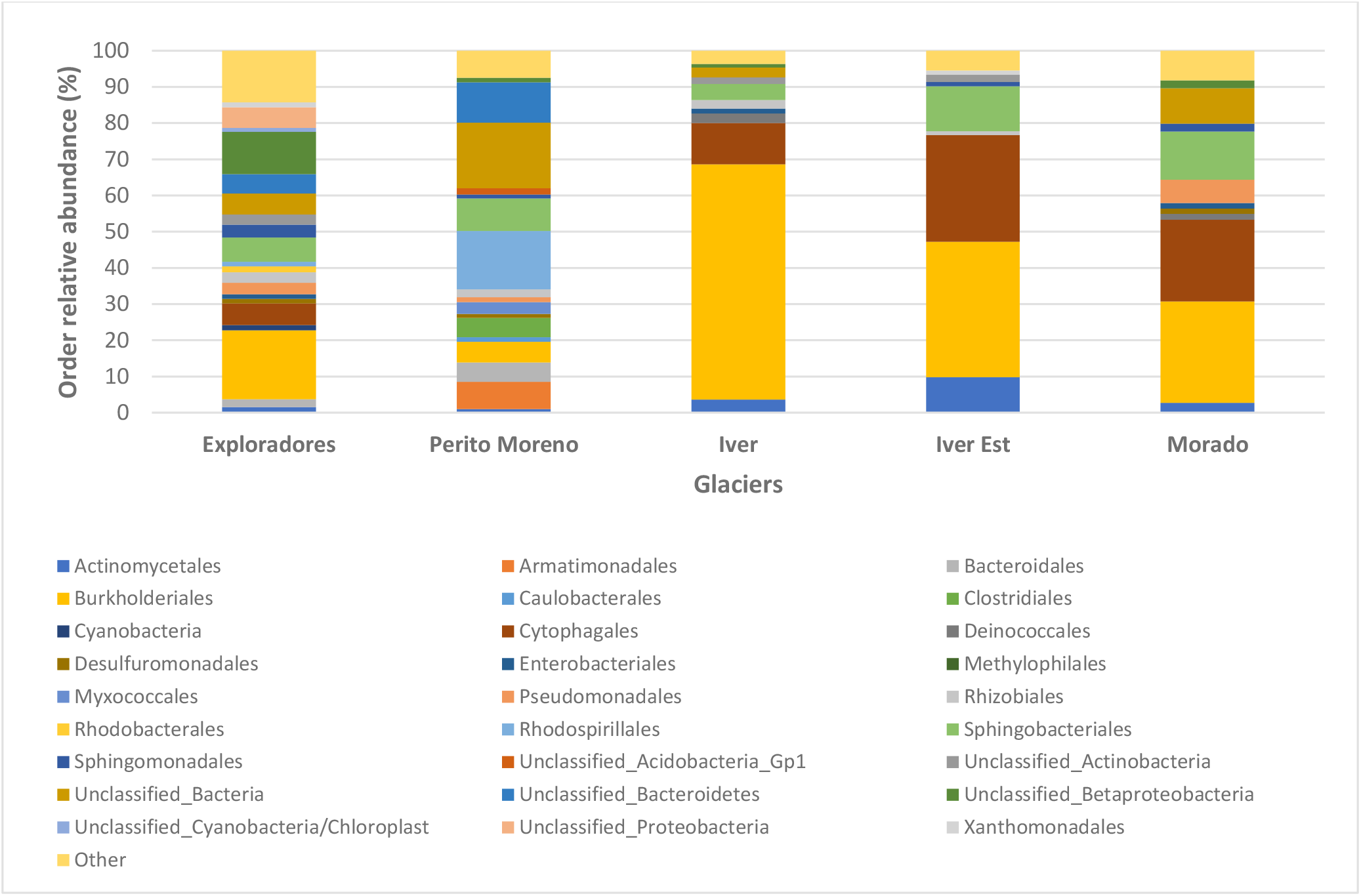
Barplot showing the relative abundance of bacterial orders in the cryoconite holes on each glacier. Orders representing less than 1 % of sequences in each sample were grouped in “others”. Cyanobacteria are reported as phylum because the rdp classifier does not provide their classification at order level.

**Figure 4.**
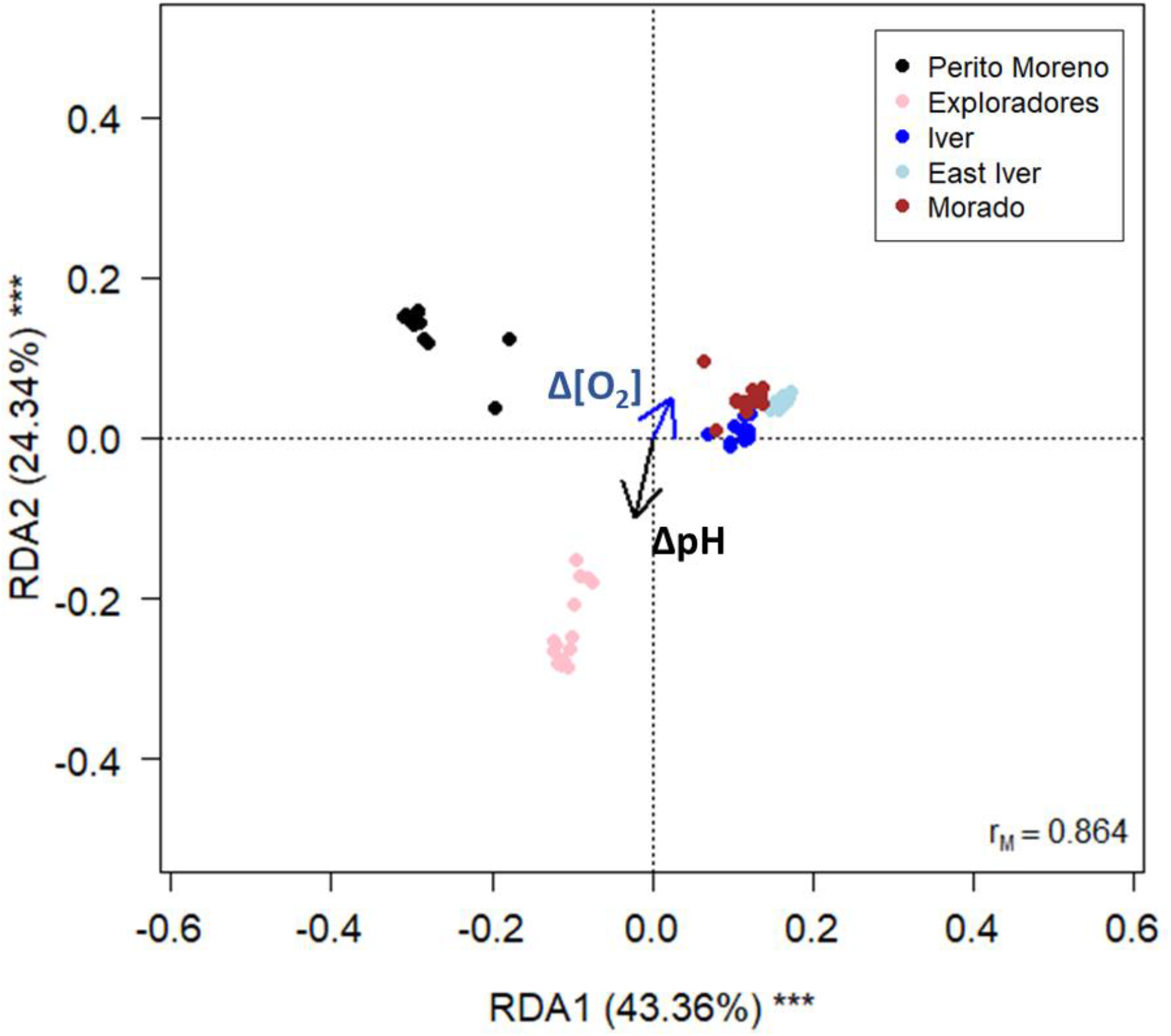
Correlation biplot from RDA on Hellinger-transformed bacterial ASVs abundances of all the samples according to glacier (five level factor), pH and oxygen concentration centred to their mean value per glacier. The blue arrow indicates increasing values of oxygen concentration in each glacier, while the black one increasing pH values in each glacier. The percentage of variance explained by each axis and their significance (***:P < 0.001) is reported. r_M_ is the Mantel correlation coefficient between the Hellinger distances between samples and the Euclidean distances between the corresponding symbols in the graph. Values close to one indicate that the graph correctly represents the distance between samples.

Post-hoc tests also revealed that the structure of the bacterial communities of all the five glaciers were significantly different from one another (|t_62_| ≥ 19.64, P_FDR_ ≤ 0.003).

GLSs showed that both the alpha diversity indexes varied significantly among glaciers (Shannon index: F_4,73_ = 42.147, P < 0.001; Gini index: F_4,73_ = 33.607, P < 0.001). Post-hoc tests showed that Shannon index was higher in Exploradores samples and lower in Iver and East Iver ones, while Gini index was higher in Iver and in East Iver samples and lower in Exploradores ones (Figure 5).

**Figure 5.**
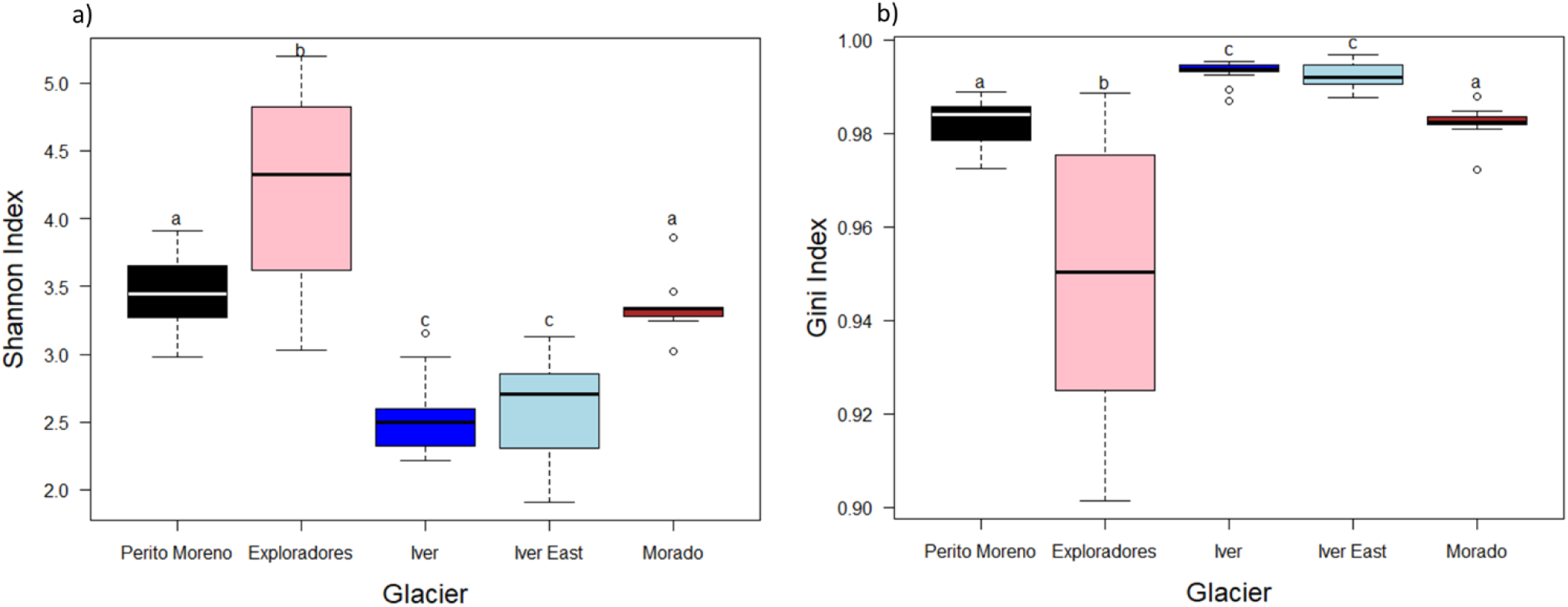
Boxplots of Shannon (a) and Gini (b) diversity indices of cryoconite hole bacterial communities. The thick lines represent the median, boxes upper and lower limits the 25^th^ and the 75^th^ percentiles respectively, whiskers the data that go beyond the 5^th^ and the 75^th^ percentile (upper whisker), dots represent the outliers and different letters indicate significant differences at post-hoc tests.

GLMs performed on the most abundant orders showed that the abundances of Burkholderiales, Cytophagales, Sphingobacteriales, Actinomycetales, Rhodospirillales, Rhizobiales, Pseudomonadales, Sphingomonadales and Bacteroidales varied among glaciers (F_4,64_ ≥ 4.207, P_FDR_ < 0.010), with complex patterns of variation. The most abundant orders varied according to the glacier without showing any particular pattern, Iver was the glacier with the highest relative abundance of Burkholderiales, Morado the glacier with the highest relative abundance of Pseudomonadales and Perito Moreno had more Rhodospirillales (Figure 6).

**Figure 6.**
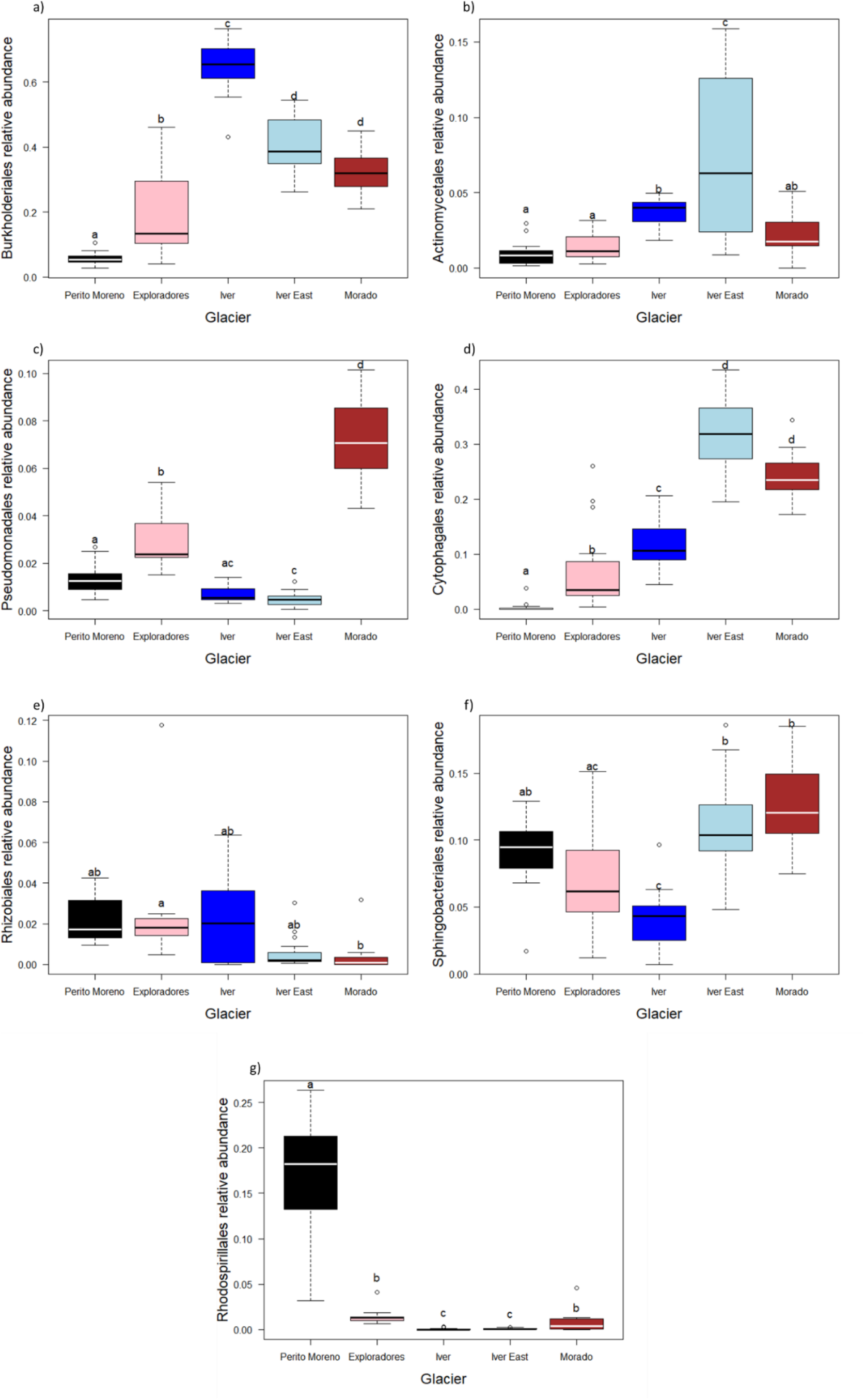
Boxplots of the relative abundances of Burkholderiales (a), Actinomycetales (b), Pseudomonadales (c), Cytophagales (d), Rhizobiales (e), Sphingobacteriales (f), Bacteroidales (g), Sphingomonadales (h) and Rhodospirillales (i) on the five glaciers where we collected cryoconite samples. The thick lines represent the median, boxes upper and lower limits the 25^th^ and the 75^th^ percentiles respectively, whiskers the data that go beyond the 5^th^ percentile (lower whisker) and the 75^th^ percentile (upper whisker), dots represent the outliers and different letters indicate differences between the mean values of different groups

## Discussion

In this study, we provide the first description of the bacterial communities of cryoconite holes from Andean glaciers, in particular from both small high elevation glaciers in the Santiago Metropolitan Region (Chile), and from the tongue of large Patagonian glaciers that reach low altitudes. Indeed, there is a big gap in our knowledge of these environments, that is the lack of data about microbial communities in supraglacial environments in this continent, that hosts about 30000 km^2^ covered by ice corresponding to about 5% of the glacierized area of the whole planet outside the Antarctica (Pfeffer et al. 2014). Results showed that the large Patagonian glaciers (Exploradores and Perito Moreno) were the two glaciers with the highest oxygen concentrations, while Iver and East Iver had the lowest ones and Morado an intermediate value. This pattern seems related to the different altitude of the glaciers. Jacobsen et al. (2003) calculated that at 4000 m a. s. l. oxygen availability in water is one fifth of that at sea level also because the kinematic viscosity of water plays an important role and depends on the elevation. This result is consistent with [O_2_] values we found in our samples. Indeed, Exploradores and Perito Moreno are located in Patagonia at low altitude (< 200 m a.s.l.), while Iver and East Iver are the highest ones among those we investigated (samples were collected at about 4000 m a.s.l.) and Morado at an intermediate value (3400 m a.s.l.).

pH values seem to follow a different pattern: the highest values were recorded on Exploradores and Morado, while the lowest were recorded on the East Iver and Iver. However, all the mean pH values of the five glaciers were basic (between 8.57 and 10.47). Differences in pH among glaciers are not easy to explain and may be due to differences in lithology of the surrounding environments that act as source of cryoconite. We note, however, that we measured water pH rather than that of the cryoconite, because the first one was already reported to influence bacterial communities in cryoconite holes (Edwards et al. 2011; Ambrosini et al. 2017). However, water pH may also be partially affected by the metabolic activities of the same bacterial communities of cryoconite holes, so it is still unclear whether water pH affects cryoconite hole bacterial communities or in part also vice versa (Hu and Cai 2011).

The most abundant orders were (in decreasing order): Burkholderiales, Cytophagales, Sphingobacteriales, Actinomycetales, Pseudomonadales, Rhodospirillales, Rhizobiales and Sphingomonadales. These orders are typical of cryoconite holes and dominate bacterial communities in these environments in all the geographical areas investigated so far: Arctic (Cameron, Hodson, and Osborn 2012; Liu et al. 2017), Antarctica (Christner, Kvitko, and Reeve 2003; Sommers et al. 2018), Europe (Edwards et al. 2013; Pittino et al. 2018), Asia (Segawa et al. 2014; Liu et al. 2017). Cryoconite holes of Andean glaciers seem therefore to host bacterial communities that are quite typical of these biodiversity hot spots on glacier environments worldwide.

Despite this general similarity of the dominant orders of bacterial communities at a global scale, a closer inspection at ASV level revealed that the glacier bacterial communities of the different glaciers differed to one another. In addition, the RDA biplot showed that the three small and high elevation glaciers clustered quite close to one another, whereas Perito Moreno and Exploradores were separate from one other and from the other glaciers. This pattern may be due to the altitude, geographic distance and ecological and environmental differences among glaciers. Indeed, bacterial communities in high elevation glaciers are exposed to high UV radiation and, therefore, to high oxidative stress (Orellana et al. 2018; Margesin and Collins 2019) and different vegetation cover (Florinsky and Kuryakova 1996). Patagonian glaciers are 440 km to one another and c.ca 2000 km from the other glaciers we sampled, which, in turn, are less than 60 km to one another. In addition, it has been demonstrated that the main source of cryoconite bacterial communities is the local environment surrounding the glacier (Stibal et al. 2015; Franzetti et al. 2017a). The high elevation glaciers we sampled are small and above the three line, and generally in areas with few vegetation. In contrast, the Patagonian glaciers where we collected are huge and their tongues (where we collected the samples) is well below the tree line and therefore exposed to very different potential sources of microorganisms with respect of the other glaciers.

The RDA also showed that ΔpH and Δ[O_2_] significantly explained the variability of bacterial communities of cryoconite holes. We stress that these variables represent the difference between pH and oxygen concentration of each cryoconite hole from the respective mean value of all the holes of that glacier. They therefore do not account for the difference in mean pH and oxygen concentration highlighted in the analyses discussed above, whose effect in this analysis is accounted for by the glacier factor, which accounts for any difference between glaciers. Thus, this analysis suggests that consistent bacterial communities vary consistently according to pH and oxygen concentration gradients present on each glacier, even if the different glaciers show on average different pH and oxygen concentration values. In other words, for example, an increase of two pH units seems to determine a similar variation in bacterial communities independently from the absolute pH value, at least in the range of variability of pH values recorded in this study. In addition, no effect of ΔpH on alpha diversity resulted from GLSs. Different studies already investigated the effect of both water and sediment pH on bacterial communities composition, proving that it influences the community in river’s sediment of different typology (i.e. glacier-fed streams, large rivers, riverine wetlands) (Shen et al. 2013; Wilhelm et al. 2013; Ligi et al. 2014; Liu et al. 2015). Nonetheless, so far the only evidence is the decrease of Acidobacteria at basic values of pH (Chu et al. 2011; Shen et al. 2013; Wilhelm et al. 2013; S. Liu et al. 2015), but in our case this effect is not evident since pH values of our samples varied from 7.25 to 12.71.

Cryoconite holes are mostly aerobic environments even in presence of an ice lid, thanks to the high O_2_ solubility in cold environments, to the release of air bubbles from the ice that melts because of the presence of the dark sediment and to photosynthesis (Zdanowski et al. 2017a; Telling et al. 2012). On these glaciers, we observed on average lower oxygen concentrations in high altitude cryoconite holes probably because water oxygen depends on the equilibrium with the atmospheric oxygen. In addition, variation in oxygen concentration seems to play an important role in explaining bacterial community structures. An effect of oxygen concentration on cryconite hole bacterial communities was already reported in previous studies on Forni glacier (Italian Alps) (Franzetti et al. 2017b). Anyway, the mechanisms that link oxygen concentration and bacterial community structure are not easy to explain, because we measured oxygen concentration in water, while a study by Poniecka et al., (2018) demonstrated that oxygen concentration into the cryoconite at the bottom of the hole is different from that in the above melting water. Indeed, the water layer works as a barrier that limits oxygen diffusion from the atmosphere to the sediment, where anoxic conditions often develop (Poniecka et al. 2018). Consistently, presence of anaerobic bacteria in cryoconite holes has been reported (Zdanowski et al. 2017a) and can be explained by the formation of anoxic niches, particularly when the sediment is thick and the water layer is deep, or within the cryoconite grains (Poniecka et al. 2018). However, measuring oxygen content in the sediment was impractical in the present study. Anyway oxygen content in the liquid phase of cryoconite holes proved to affect bacterial communities inhabiting cryoconite, even if we know that this sediment is anoxic after few micrometers of depth (Poniecka et al. 2018). Nevertheless, it was already observed an effect of dissolved oxygen concentration on bacterial communities of the intertidal biofilm in estuarine waters (Guo et al. 2017), and that bacterial communities change along a vertical profile in river’s sediment because of the switch from oxic to anoxic conditions (Huang et al. 2011). In marine sediment, it was observed that going deeper in the sediment, where dissolved oxygen decreased, diversity of nitrifying bacteria decreased probably because they are facultative aerobic and in the shallowest part they have mostly an aerobic metabolism, and also because of the deposition and the consequent decomposition and transformation of organic matter that provides more nutrients availability (Tiquia, Masson, and Devol 2006). Unfortunately, in our case, the sediment depth is very limited (mostly < 0.5 cm) and it is not possible to obtain the vertical distribution of bacterial taxa along the vertical profile of the sediment, like in rivers and marine sediment where the effect of oxygen absence was visible after half centimetre of few centimetres (Tiquia, Masson, and Devol 2006; Huang et al. 2011). Anyway, this result proved that, even if mostly anoxic, the cryoconite layer microbial community is affected by the oxygen dissolved in water. Its direct affect likely acts on the intertidal bacterial biofilm, that represents a small fraction of the whole community, so probably the effect is appreciable at the whole community level because of the interactions that subsist between different *taxa*, and going deeper in the cryoconite layer we may see an indirect effect of dissolved oxygen.

The glaciers Iver and East Iver are the two geographically closest glaciers (only 56 km apart), and they were also the most similar ones in oxygen concentration and pH (Fig. 2a-b) as well as in alpha diversity values and in the structure of bacterial communites, as shown by the fact that they were close to one another in the RDA biplot. Interestingly, also the Morado glacier clustered close to them in the RDA biplot, even if it is c.ca 60 km from them. In contrast, the two Patagonian glaciers are very far on the same plot. These results, on the one side, support the hypothesis that the altitude and the geographic position plays an important role in defining bacterial community composition. On the other side, however, the three small glaciers in central Andes are also at similar elevation and they are surrounded by very similar environments (R. Ambrosini and F. Pittino, personal observation). Their similarity can therefore derive also from being exposed to the same general ecological conditions including high UV radiation and oxidative stress and probably to similar sources of bacterial communities. These results therefore highlight that correlative studies like the present ones can hardly disentangle the effects of geographical positions and ecological conditions on the structure of cryoconite hole bacterial communities, and further studies should be designed to add insight into this still open question.

Analyses on alpha diversity indicate that cryoconite holes on Exploradores glacier showed the highest richness and evenness. This may be because Exploradores is located in an area surrounded by a mixed broadleaf forests, whose canopy should host rich and diverse microbiome, which can act as a source of bacterial communities. The other Patagonian glacier, Perito Moreno, despite being below the treeline in the sampled area, is surrounded by a less diverse vegetation (woods dominated by southern beeches, *Nothofagus* ssp.), which can be a less diverse source of bacteria. Indeed, the analyses showed that the alpha diversity values of cryoconite holes of this glacier did not differ significantly from those of small glaciers in central Andes, which were above the treeline. In addition, samples on the Exploradores were collected close to the glacier terminus and in an area with abundant supraglacial debris and frequented by tourists. In contrast, Perito Moreno samples were collected farther from the glacier border and in an area with less sparse supraglacial debris, visibly cleaner and not frequented by tourists. Therefore, the surrounding forest did not affect cryoconite bacterial communities as much as on the Exploradores glacier, where samples were collected in an area closer to the glacier terminus and to the surrounding forest. Of course, sampling a glacier in a more exhaustive way collecting samples from almost the whole area of the glacier would give a more complete overview of the communities present there, but for vast glaciers (like Perito Moreno and Exploradores) it would be very complicated.

It looks like the main differences may be due to the belonging to a high elevation rather than a low elevation glacier. Looking at the results about differences of the most abundant orders, Cytophagales, Burkholderiales and Actinomycetales are the three orders that show different abundances according to the belonging to low or high elevation glaciers (Fig. 7a-b-d). Burkholderiales are a quite heterogeneous order, and therefore it is difficult to understand their trend according to an ecological interpretation (Garrity, Bell, and Lilburn 2015). Cytophagales are gram negative bacteria (Reichenbach 2006), that are known to be less resistant to UV radiation than gram positive bacteria (Arrage et al. 1993), therefore it is unexpected their higher relative abundance in bacterial communities of Iver, Iver East and Morado glacier. The order Actinomycetales, on the other hand, are mostly Gram positive bacteria (Cummins and Harris 1958), that is consistent with their higher relative abundance in high elevation glaciers since they are more resistant to high UV radiation (Arrage et al. 1993).

Bacterial communities of cryoconite holes show seasonal variation on temperate mountain glaciers (Pittino et al. 2018), therefore it may be argued that the differences we observed in the community structures among glaciers may be due, at least partly, to differences in the stages of the seasonal ecological succession present on each glacier when we collected the samples. In other words, one may argue that the bacterial communities were identical on each glacier at the beginning of the melting season and changed seasonally according to identical ecological succession, but they appear different in our samples because we collected them at different stages. Despite we acknowledge that this process may contribute to the observed variability among glaciers, we consider this effect as minor, and we confidently suggest that the differences we observed depend mostly on spatial variation and on variation in the general ecological conditions of the glacier and of the surrounding environments (mostly altitude and vegetation). Indeed, even if microbial communities can change along the ablation season, there always is a core community characteristic of one glacier. Furthermore the differences among glaciers we see in our samples’ bacterial communities, are not ascribable to a temporal trend like the one described by Pittino et al. (2018). Indeed, we do not see any difference in phototrophs and heterotrophs abundances, and Cyanobacteria had a relative abundance higher than 1 % only on the Exploradores glacier. While if there was a temporal trend, Cyanobacteria should have been more present on those glaciers where the ablation season started later, but this is not the case.

In summary we provide the first-ever description of the bacterial communities of cryoconite holes of glaciers in South America, which confirm that these environments are dominated by the same bacterial orders all over the world (Boetius et al. 2015; Liu et al. 2017; Christner, Kvitko, and Reeve 2003; Franzetti et al. 2016). The dissolved oxygen concentration in the water seems to affect bacterial communities that are mostly anoxic. Water pH also influenced bacterial communities. Importantly, this study, is not based on one glacier only, and therefore, can also give some insights on the ecological features that drive the structure of cryoconite hole bacterial communities on different glaciers. The five glaciers we investigated are still a too small sample for thoroughly assess the ecological processes that control cryoconite hole bacterial communities, but this is still a much larger sample size than that on which the vast majority of the studies on cryoconite holes published so far was based and it can put the basis to further investigations aiming at understanding how different and, at the same time, how similar cryoconite holes bacterial communities are.

## Notes

### Competing Interest Statement

The authors have declared no competing interest.

